# Functional Assessment and Extended Culture of Human Pediatric Cardiac Tissue Slices for Preclinical Testing

**DOI:** 10.64898/2026.11.04.762282

**Authors:** Devon Guerrelli, Kathleen McLean, Allison Freeman, Jenna Pressman, Alisa Bruce, Manan Desai, Yves d’Udekem, Nikki Gillum Posnack

**Author notes:** Corresponding Author: Nikki Gillum Posnack, Ph.D., Sheikh Zayed Institute for Pediatric Surgical Innovation Children’s National Hospital, 111 Michigan Avenue, NW Washington, DC, USA 20010. Authors contributed equally to this work (Kathleen McLean, Allison Freeman).

## Abstract

**Background:** Living human myocardial slices preserve native tissue architecture, but their application to pediatric cardiac physiology or prolonged functional testing remains limited. We adapted a passive culture method for small surgical myocardial specimens obtained from patients with congenital heart disease (CHD).

**Methods:** Atrial and ventricular specimens were obtained from 41 cardiac surgery patients (7.7±11.1 years; 1 day – 49 years). Myocardial tissue was sectioned into slices and studied acutely, or maintained under free-floating, rocking cell culture conditions without electrical or mechanical stimulation. Calcium transients were recorded in response to external pacing (0.2–2 Hz) at baseline, and after isoproterenol or norepinephrine exposure. Selected preparations underwent viability, histological, and gene-expression analyses.

**Results:** Pacing-evoked calcium transients were recorded from all four cardiac chambers, exhibiting expected frequency-dependent shortening. Mean interslice and intraslice coefficients of variation were 11.1% and 7.6%. Viability was 85.8 ± 5.6% post-slicing and 78.8 ± 5.0% after 3–5 hours. Calcium transients remained detectable through 178 days in atrial and 119 days in ventricular slices. Calcium transient durations were relatively stable during extended culture. Beta-adrenergic agonists shortened calcium transient durations (isoproterenol: 32% atrial, 25% ventricular at 0.3 Hz); responses to isoproterenol persisted to approximately 120 days in selected atrial slices and 60 days in ventricular slices. Selected histological features were preserved, although some extracellular-matrix and apoptosis-related transcripts changed.

**Conclusions:** Small surgical specimens support repeated physiological measurements and pharmacological testing during extended passive culture, providing a platform for human developmental and disease-specific cardiac research.

## INTRODUCTION

Congenital heart disease (CHD) is the most common birth defect, affecting approximately 1% live births each year(Van Der Linde et al., 2011). Advances in surgical and medical care have substantially improved survival, as 97% of babies born with CHD now survive into adulthood(Mandalenakis et al., 2020). Nevertheless, many of these patients will require repeated clinical interventions and/or pharmacological therapies throughout their childhood. Indeed, cardiovascular medications are frequently administered to pediatric CHD patients off-label(Pasquali et al., 2008), with limited information regarding the safety and efficacy on the developing heart. Experimental models that preserve native pediatric myocardial properties are therefore needed to understand developmental physiology, disease-associated remodeling, and drug responsiveness.

The developing myocardium undergoes extensive molecular, structural, and physiological changes after birth – including changes in ion channel expression, excitation-contraction coupling, intracellular calcium handling, and mechanical function(Salameh et al., 2023). Although studies on isolated human pediatric cardiomyocytes or myocardial muscle strips are limited, published work has identified important functional differences across development. For example, Wiegerinck et al. demonstrated that the force-frequency relationship is relatively flat in the newborn human ventricular myocardium but becomes more positive during infancy, coinciding with changes in sodium-calcium exchanger and phospholamban expression(Wiegerinck et al., 2009). Elhamine et al. further identified developmental changes in myofibrillar protein isoforms, calcium sensitivity, and contractile kinetics in pediatric ventricular tissue(Elhamine et al., 2016). More recently, our laboratory identified extensive age-dependent changes in human atrial gene expression, including ion channels, calcium-handling proteins, and contractile machinery(Salameh et al., 2024). These findings demonstrate that pediatric myocardial physiology differs substantially across developmental stages, but functional studies using intact human tissue remain limited.

Currently, preclinical approaches to studying cardiac developmental physiology rely heavily on animal models and human induced pluripotent stem cell-derived cardiomyocytes (hiPSC-CM). Although hiPSC-CMs support scalable pharmacological screening, their developmental immaturity limits direct comparisons with native pediatric myocardium(Guo & Pu, 2020; Karbassi et al., 2020). Indeed, transcriptomic analyses have demonstrated that hiPSC-CMs closely resemble first-trimester human fetal myocardium, with maturation-promoting culture conditions shifting expression profiles toward second-trimester fetal stages(Van Den Berg et al., 2015).

Animal models preserve integrated cardiac electrical and mechanical function, but species-dependent differences can limit clinical translation. For example, mice and rats have substantially higher intrinsic heart rates, shorter ventricular action potentials with limited plateau phases, and greater dependence on sarcoplasmic reticulum calcium cycling than humans(Blackwell et al., 2022; Ripplinger et al., 2022). Differential protein expression and calcium-handling mechanisms also contribute to species-specific contractile and relaxation kinetics(Janssen et al., 2016; Linscheid et al., 2021; Milani-Nejad & Janssen, 2014). Although rabbits, guinea pigs, and larger mammals more closely approximate aspects of human cardiac electrophysiology, important differences in ionic currents, tissue organization, and physiological regulation remain(Odening et al., 2021; Ripplinger et al., 2022). Collectively, these limitations highlight the need for complementary human tissue models that retain native myocardial properties and permit direct assessment of developmental and disease-associated differences in cardiac function.

One such emerging technology is precision-cut living myocardial slices, often referred to as “organotypic slices,” which preserve the native three-dimensional architecture and multicellular organization of cardiac tissue. By reducing tissue thickness to a few hundred micrometers, this approach facilitates oxygen and nutrient diffusion from the surrounding culture medium, supporting tissue viability and physiological function(Van Der Geest et al., 2025; Watson et al., 2020). Prior studies have established the feasibility of using human myocardial slices for electrophysiology, calcium handling, contractility, and drug safety assessments(Brandenburger et al., 2012; Fischer et al., 2019; Kang et al., 2016; Ou et al., 2019; Watson et al., 2017). However, these studies have relied predominantly on adult myocardial tissue, typically obtained from donor hearts unsuitable for transplantation or diseased hearts explanted during cardiac transplantation. Consequently, the application of this technology to pediatric CHD myocardium remains limited. Pediatric surgical specimens also present additional challenges, as they are often small, anatomically heterogeneous, and available only when tissue is resected during clinically indicated cardiac surgery.

In this study, we adapted previously established cardiac slice-culture methods incorporating continuous medium agitation(Fischer et al., 2019; George et al., 2020; Kang et al., 2016; Qiao et al., 2019) to maintain viable myocardial tissue slices from surgically resected tissue that would otherwise be discarded during cardiac surgery. We assessed baseline calcium cycling across all four chambers, frequency-dependent calcium restitution, intra-and interslice variability. We also evaluated changes in tissue architecture and selected molecular markers during extended culture, along with beta-adrenergic responsiveness to isoproterenol and norepinephrine. These studies establish a practical platform for prolonged physiological and pharmacological assessment of human pediatric myocardium, thus maximizing the experimental utility of valuable surgical specimens.

## METHODS

### Clinical Specimen and Data Collection

This study was approved by the Children’s National Hospital Institutional Review Board (IRB Pro00012146 / STUDY00000198), and all experiments followed applicable human research guidelines. Patients undergoing clinically-indicated cardiac surgery were approached for participation. Following surgical resection, tissue specimens were transported to the research laboratory. Patient demographic data was retrospectively extracted from electronic medical records, including patient age, sex, diagnosis, and surgical procedure (**Table 1**).

**Table 1:** Patient & Tissue Sample Characteristics. ATR SEP = atrial septum, ASD = atrial septal defect, AVSD = atrioventricular septal defect, DILV = double inlet left ventricle, F = female, HLHS = hypoplastic left heart syndrome, IVS = intact ventricular septum, LAA = left atrial appendage, LV = left ventricle, M = male, MAPCAs = major aortopulmonary collateral arteries, PAPVC = partial anomalous pulmonary venous connection, PAPVR = partial anomalous pulmonary venous return, RA = right atria, RAA = right atrial appendage, RV = right ventricle, TGA = transposition of the great arteries, VSD = ventricular septal defect. n/a = not applicable; slices imaged acutely and saved for alternate applications (RNA analysis & histology). Slices still alive at the time of publication are denoted with a > symbol in the culture duration column.

| Age | Age Group | Sex | Fundamental Diagnosis | Sample Type | Culture Duration (days) |
|---|---|---|---|---|---|
| 1 day | Neonate | M | Ebstein's anomaly | RA | 120 |
| 2 days | Neonate | M | Tricuspid valve disease | RA | 108 |
| 11 days | Neonate | F | Tricuspid valve disease, Ebstein's anomaly | RA | n/a |
| 15 days | Neonate | M | HLHS | RV | 28 |
| 27 days | Neonate | F | Mitral valve disease, Mitral regurgitation | RA | 119 |
| 29 days | Neonate | F | Ebstein's anomaly | RA | 178 |
| 56 days | Infant | M | LV aneurysm, Depressed LV function | LV | >28 |
| 107 days | Infant | M | Tetralogy of Fallot, Pulmonary stenosis | RV | 69 |
| 112 days | Infant | F | Tetralogy of Fallot, Pulmonary stenosis | RV | 48 |
| 113 days | Infant | M | Pulmonary atresia, VSD | RV | 99 |
| 121 days | Infant | M | Tetralogy of Fallot, Pulmonary stenosis | RV | 48 |
| 134 days | Infant | M | Pulmonary atresia, IVS | ATR SEP | 28 |
| 147 days | Infant | F | Mitral valve disease, Mitral regurgitation | RA | 90 |
| 197 days | Infant | M | Pulmonary atresia, VSD, MAPCAs | RAA | 110 |
| 214 days | Infant | M | Congenitally corrected TGA, VSD | RV | 90 |
| 224 days | Infant | F | Transitional AVSD | RAA | n/a |
| 278 days | Infant | M | Single ventricle, DILV | ATR SEP | n/a |
| 2.6 years | Child | M | Cardiomyopathy (Hypertrophic, Obstructive) | LV | 118 |
| 2.7 years | Child | F | ASD | RAA | n/a |
| 3.3 years | Child | M | VSD | RV | 119 |
| 3.8 years | Child | F | Pulmonary atresia, VSD | RA | n/a |
| 3.9 years | Child | M | ASD, PAPVR | RA | 60 |
| 4.3 years | Child | F | Truncus arteriosus, Interrupted aortic arch | RV | 28 |
| 4.8 years | Child | M | Single ventricle, Tricuspid atresia | ATR SEP | 120 |
| 5.2 years | Child | M | HLHS | LV | 60 |
| 6.0 years | Child | F | Aortic stenosis | LV | 120 |
| 6.1 years | Child | F | Complete AVSD | LV | 28 |
| 8.8 years | Child | M | ASD, PAPVR | RAA | n/a |
| 9.3 years | Child | F | Pulmonary atresia, VSD | RV | 19 |
| 10.4 years | Child | F | Truncus arteriosus | RV | 29 |
| 10.5 years | Child | F | ASD, PAPVR | RAA | >28 |
| 11.1 years | Child | F | PAPVC | RAA | 150 |
| 12.5 years | Child | M | Complete AVSD | LV | 22 |
| 13.3 years | Child | M | Cardiomyopathy (Hypertrophic, Obstructive) | LV | n/a |
| 15.7 years | Child | M | ASD | RAA | n/a |
| 17.6 years | Child | F | Coronary anomaly | RAA | 90 |
| 18.0 years | Adult | M | Cardiomyopathy (Hypertrophic) | LV | 70 |
| 20.5 years | Adult | M | Aortic stenosis | RV | n/a |
| 31.3 years | Adult | M | Aortic stenosis | RV | 29 |
| 40.8 years | Adult | F | ASD | RV | 78 |
| 49.1 years | Adult | M | Coarctation of the Aorta | LAA | 118 |

### Preparation of Live Myocardial Tissue Slices

An overview of the protocol is summarized in **Figure 1**. Heart specimens were transported from the operating room to the laboratory. Specimens were trimmed and then secured to a standard tissue mount (7000-4-1; Campden Instruments) using Dermabond (DHVM12). The mounted samples were then submerged in a tissue bath containing cold Tyrode’s slicing solution(Watson et al., 2017) (30 mM 2,3-butanedione monoxime (BDM), 1 mM glucose, 10 mM HEPES, 6 mM potassium chloride, 140 mM sodium chloride, 1 mM magnesium chloride, and 1.8 mM calcium chloride; **Supplemental Table S1**), bubbled with 100% oxygen. A higher concentration of potassium and BDM were included to minimize spontaneous electrical activity and contraction during the slicing procedure. An external ice bath was used to maintain the slicing temperature near 4°C. In accordance with established protocols(Kang et al., 2016; Qiao et al., 2019), a high-precision vibratome (7000smz-2; Campden Instruments) with a stainless-steel blade was used to cut tissue samples into 380 μm slices using 0.04 mm/s advance speed, 2 mm horizontal vibration amplitude, 80 Hz vibration frequency, and a z-axis vibration of less than 0.5 µm. Pediatric myocardial specimens that were too small to secure on the vibratome stage were manually dissected into small tissue fragments(Usta et al., 2010). Cardiac tissue slices were immediately transferred to a room temperature Tyrode’s recovery solution (1 mM glucose, 10 mM HEPES, 4.5 mM potassium chloride, 140 mM sodium chloride, 1 mM magnesium chloride, and 1.8 mM calcium chloride; **Supplemental Table S2**), bubbled with 100% oxygen. During recovery, 100 μm nylon mesh strainers were used to support cardiac tissue slices at the liquid-air interface; slices recovered for 20 min to permit equilibration to room temperature and to wash out BDM. Cardiac tissue slices were subsequently transferred to a tissue culture incubator or used immediately for experimentation.

**Figure 1.**
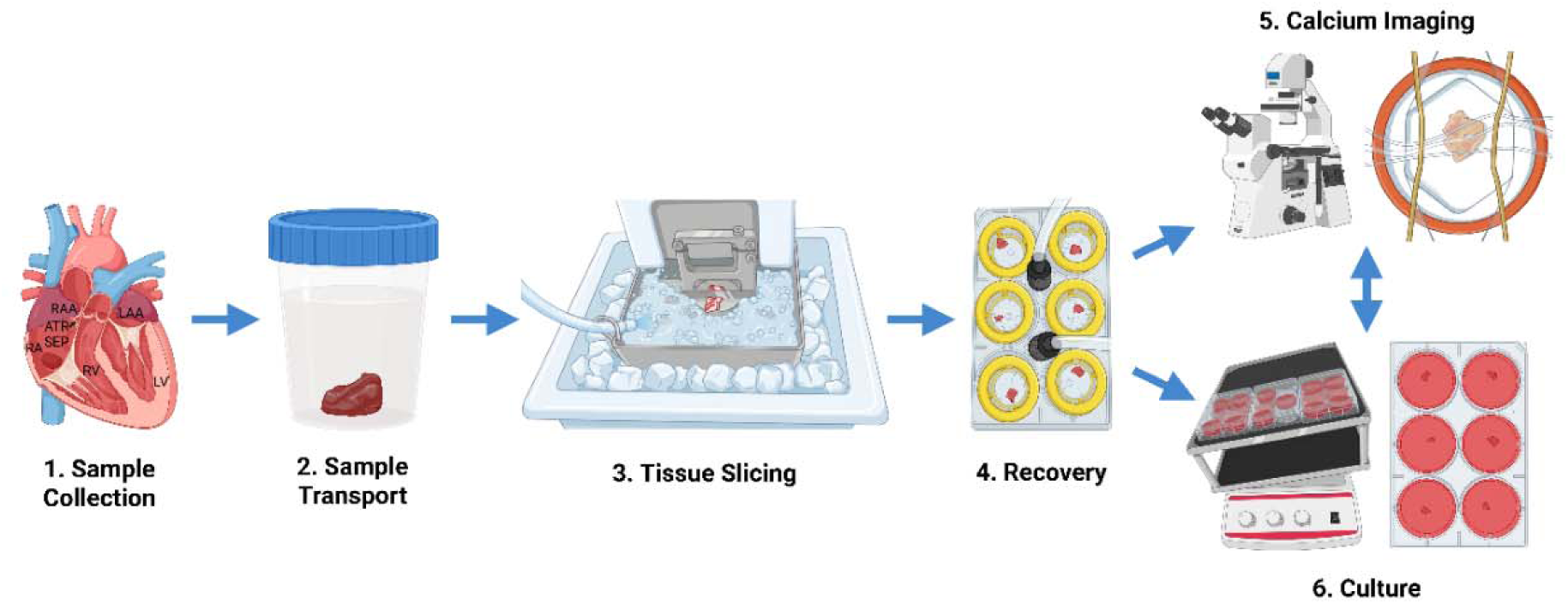
Overview of human myocardial tissue slice preparation and culture. **1)** Collection of small myocardial specimens that would otherwise be discarded during cardiac surgery and **2)** rapid transport to the research laboratory. **3)** Tissue specimens are sectioned into 380-µm slices using a vibratome in ice-cold, oxygenated Tyrode’s slicing solution. **4)** Slices are transferred to Tyrode’s recovery solution for _≥_20 minutes at room temperature. **5)** Following recovery, slices are used for intracellular calcium imaging or **6)** maintained under passive culture conditions for subsequent functional assessments.

### Passive Culture of Human Myocardial Slices

Tissue slices were transferred to individual wells of a six-well plate, and remained free-floating in 7 ml of M199 culture media supplemented with 1x ITS liquid media supplement, 40 U/ml penicillin, and 40 μg/ml streptomycin (**Supplemental Table S3**). Tissue slices were cultured in a standard cell culture incubator at 37°C with 5% CO_2_. During culture, slices were rocked on a rocking shaker (SHRK07AL2, Ohaus) with a speed of 4 and a tilt of 6. Media was replaced every 3-4 days. For extended culture, tissue slices were transferred to a new six-well plate every 2 weeks. For repeat experiments, approximately 10 days elapsed between imaging sessions.

### Live Fluorescent Imaging

Tissue slices were loaded with a calcium indicator dye (20 µM Fluo-4AM, ThermoFisher Scientific) in the dark at room temperature for approximately 40 min, and then washed in Tyrode’s Recovery Solution. For electrical stimulation, tissue slices were positioned into an imaging chamber with field stimulation (RC-49MFSH, Warner Instruments) that was modified with fishing line to gently secure the slice in place. Intracellular calcium transients were recorded using an inverted Nikon microscope (475/525 nm excitation/emission) equipped with a high-speed camera (Prime 95B, Photometrix). Calcium transients were recorded at room temperature in response to external stimulation (0.2 – 2 Hz) at baseline and after acute 5 min exposure to 1 μM norepinephrine or isoproterenol. In a subset of studies, tissue slices underwent repeat isoproterenol challenges during extended culture (<1 week, 2 months, 4 months). We measured calcium transient duration (CaD) to 30%, 50%, and 80% recovery (CaD30, CaD50, CaD80) using a custom MATLAB script, averaging at least 5 transients per recording.

### Tissue Viability and Histology

Tissue viability was assessed 30 min and 3-5 hours after slicing. Tissue slices were stained with a ReadyProbes Cell Viability Imaging Kit (R37610; ThermoFisher Scientific) for 15 minutes in the dark at room temperature. The number of live (NucBlue) vs dead cells (propidium iodide) was quantified using a custom MATLAB script. For histology, tissue slices were fixed in 10% formalin, paraffin embedded, and sliced to 4 µm thickness. Specimens were rehydrated through a graded ethanol procedure and antigen retrieval was performed using a citrate buffer (C999, Millipore Sigma) in a pressure cooker. Primary antibodies included α-actinin (701914; ThermoFisher Scientific) and N-cadherin (PA5-19486; ThermoFisher Scientific) at a dilution of 1:200 overnight at 4°C; goat anti-rabbit Alexa Fluor 488 (A32731; ThermoFisher Scientific) was used as a secondary antibody at a dilution of 1:500 for 1 hour at room temperature. Slides were mounted with Fluoromount-G with DAPI (00-4959-52; ThermoFisher Scientific), which was used as a nuclear counter stain. Fluorescence imaging was performed on a Leica TCS SP8 confocal microscope. Hematoxylin and eosin stains were used for histological assessment of nuclei, cytoplasm, and extracellular matrix; slides were imaged with a Lionheart FX microscope. Two blinded observers (AF, KM) scored myofiber alignment, interstitial space, nuclear morphology, and myocyte density on a 0-4 scale to assess gross structural changes in tissue slices during extended cell culture.

### Apoptotic & Fibrotic Gene Expression Analysis

Gene expression was assessed as previously described(Salameh et al., 2024, 2026). Briefly, cardiac tissue slices were submerged in RNAlater (AM7020; Invitrogen) and stored at 4°C for >24 hours. RNAlater was then removed and specimens were stored at -80°C until RNA isolation. A RNeasy Fibrous Tissue Mini Kit (74704; Qiagen) was used for RNA isolation; RNA quantity and quality were assessed using a NanoDrop spectrophotometer or Qubit instrument. Clariom S microarrays (902927; ThermoFisher Scientific) were used for transcriptomics analysis of tissue slice samples collected during extended culture. Gene expression values were z-score normalized separately for each gene across all tissue slices. Candidate genes associated with extracellular matrix remodeling and apoptotic signaling were selected based on prior myocardial slice studies(Fischer et al., 2019; Nunez-Toldra et al., 2022; van der Geest et al., 2025; Waleczek et al., 2025), with Gene Ontology annotations reviewed using Enrichr (GO:0032964, 1900117)(Chen et al., 2013; Kuleshov et al., 2016).

### Data Reporting & Statistical Analysis

Data are reported as mean ± standard deviation (SD), unless otherwise indicated. Percent changes are expressed relative to baseline measurements. Atrial data are shown in red and ventricular data in blue. Two-group comparisons were performed using two-tailed Student’s t-tests, with paired tests used for repeated measurements from the same tissue slice or patient. Frequency-dependent changes in calcium transient duration were assessed using two-way ANOVA, with pacing frequency and calcium transient duration (CaD30, CaD50, CaD80) as factors. Longitudinal changes in calcium transient duration during extended culture were assessed using one-way ANOVA, with comparisons relative to early culture (days 0–5) or days 8–10. Selected gene-expression and histological measurements were analyzed using ANOVA with culture time as the primary factor. Gene-expression comparisons were performed relative to days 8–10, whereas histological comparisons used days 0–1 or days 28–30 as reference periods. Holm–Šidák correction was applied for multiple comparisons, as appropriate. Statistical significance was defined as p < 0.05.

## RESULTS

### Preparation of Functional Human Myocardial Slices from Surgical Specimens

We adapted established cardiac tissue slicing methods to generate living myocardial preparations from surgically resected tissue that would otherwise be discarded (**Figure 1**). Following surgical resection, specimens were transported to our laboratory, secured to a tissue mount, and sectioned using a vibratome. Tissue slices were allowed to recover for at least 20 minutes before immediate experimentation or transfer to culture for subsequent studies. Live/dead staining in a sample subset demonstrated 85.8 ± 5.6% viable cells immediately following tissue preparation and 78.8 ± 5.0% after 3–5 hours (p<0.05; **Supplemental Figure S1**). Representative histological and immunofluorescence images demonstrated recognizable myocardial architecture, including sarcomeric α-actinin and N-cadherin localization post-slicing (**Supplemental Figure S1**). We subsequently evaluated myocardial specimens from N=41 cardiac surgery patients ranging in age from 1 day to 49 years, including 6 neonates, 11 infants, 19 children, and 5 adults (**Table 1**). Specimens originated from multiple anatomical regions, including 20 atrial samples (16 right atrial, 3 atrial septal, and 1 left atrial) and 21 ventricular samples (13 right ventricular and 8 left ventricular). These specimens were used to establish a workflow for functional assessments of human myocardium across developmental ages and cardiac chambers.

### Frequency-Dependent Calcium Cycling Across All Four Cardiac Chambers

We successfully recorded intracellular calcium transients from myocardial preparations representing all four chambers of the heart (**Figure 2A**), including the right atrial appendage (RAA), right atrial free wall (RA), atrial septum (ATR SEP), left atrial appendage (LAA), right ventricle (RV), and left ventricle (LV). Field stimulation at frequencies ranging from 0.2 to 2.0 Hz elicited calcium transients, allowing assessment of frequency-dependent calcium cycling (**Figure 2B–E**). Myocardial preparations from all four chambers exhibited the expected frequency-dependent changes in calcium transient kinetics, with progressively shorter durations at faster pacing rates.

**Figure 2:**
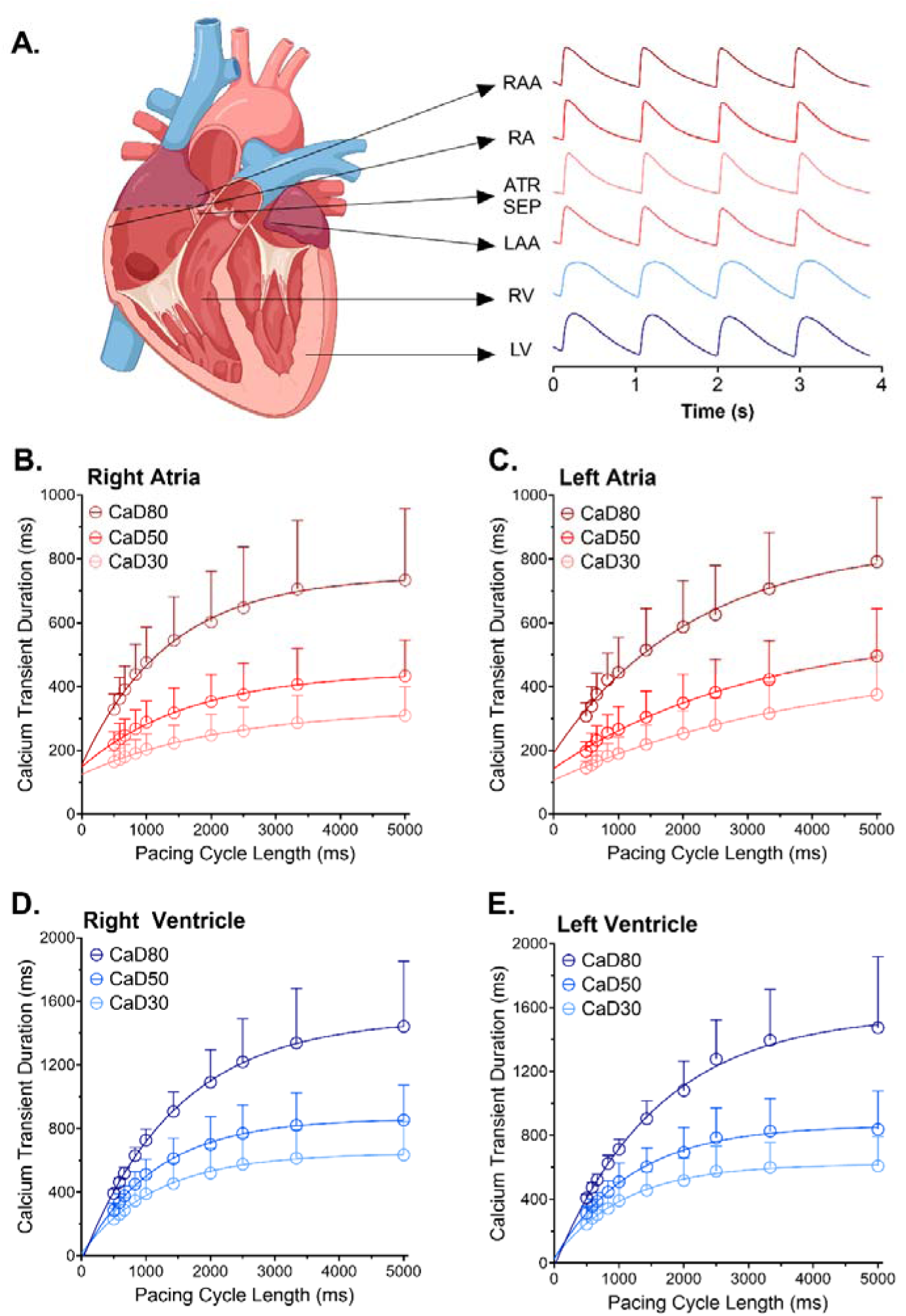
Frequency-dependent calcium cycling in human myocardial slices from all four cardiac chambers. **A)** Representative intracellular calcium transient recordings from six anatomical regions at 1 Hz pacing frequency: right atrial appendage (RAA), right atria (RA), atrial septum (ATR SEP), left atrial appendage (LAA), right ventricle (RV), and left ventricle (LV). **(B–E)** Calcium transient restitution curves showing calcium transient duration at 30% (CaD30), 50% (CaD50), and 80% (CaD80) recovery as a function of pacing cycle length (ms) in the **(B)** right atrium, **(C)** left atrium, **(D)** right ventricle, and **(E)** left ventricle. Recordings were acquired at room temperature during electrical pacing (0.2–2.0 Hz). Right atrial recordings include both appendage and free-wall preparations. Sample sizes and patient ages were as follows: right atrium (n=44 slices, N=9 patients, 1 day–17 years); left atrium (n=13 slices, N=1 patient, 49 years); right ventricle (n=25 slices, N=8 patients, 2 weeks–40 years); and left ventricle (n=30 slices, N=5 patients, 2–18 years). Data are presented as mean + SD.

A stepwise increase in pacing frequency from 0.2 to 2.0 Hz significantly shortened the mean calcium transient duration at 80% recovery (CaD80) by 55.3% in the right atrium, 61.2% in the left atrium, 72.8% in the right ventricle, and 72.3% in the left ventricle (% change, p<0.0001 for all chambers). At 0.5 Hz, CaD80 was 602.1 ± 159.4 ms in the right atrium, 587.8 ± 144.3 ms in the left atrium, 1091.4 ± 204.8 ms in the right ventricle, and 1081.5 ± 184.1 ms in the left ventricle. At 1 Hz, CaD80 shortened to 475.5 ± 110.5 ms in the right atrium, 444.6 ± 110.1 ms in the left atrium, 726.9 ± 69.1 ms in the right ventricle, and 713.8 ± 61.6 ms in the left ventricle (p<0.0001 for 0.5 vs 1.0 Hz in all four chambers). Similar frequency-dependent responses were observed for CaD30 and CaD50. Of note, right atrial measurements included slices from both the atrial appendage and free wall. Calcium imaging was performed at room temperature. Under these conditions, atrial preparations exhibited shorter calcium transient durations than ventricular preparations, consistent with previously reported chamber-specific differences in human myocardial calcium cycling(Kang et al., 2016; Maier et al., 2000). Together, these findings demonstrate that small surgical myocardial specimens can yield functional preparations capable of generating measurable, pacing-dependent intracellular calcium transients across all four cardiac chambers.

### Intra-and Interslice Variability in Calcium Transient Measurements

Prior studies have demonstrated regional and transmural heterogeneity in calcium handling within myocardial tissue(Lou et al., 2011; Wen et al., 2018). Furthermore, anatomical differences between sequential myocardial slices may contribute to variability in physiological measurements(Van Der Geest et al., 2025). To assess the consistency of our calcium measurements, we first quantified interslice variability using 5–10 slices each collected from six individual patient samples (**Figure 3A,B**). The coefficient of variation (CV) ranged from 7.8– 17.8% for CaD30, 6.8–14.1% for CaD50, and 6.1–10.9% for CaD80, with an overall mean CV of 11.1% across measurements. We also assessed intraslice variability by recording calcium transients from 5 distinct regions within individual atrial and ventricular slices (**Figure 3C,D**). The CV ranged from 2.6–12.0% for CaD30, 1.9–12.8% for CaD50, and 1.9–12.2% for CaD80, with an overall mean CV of 7.6%.

**Figure 3.**
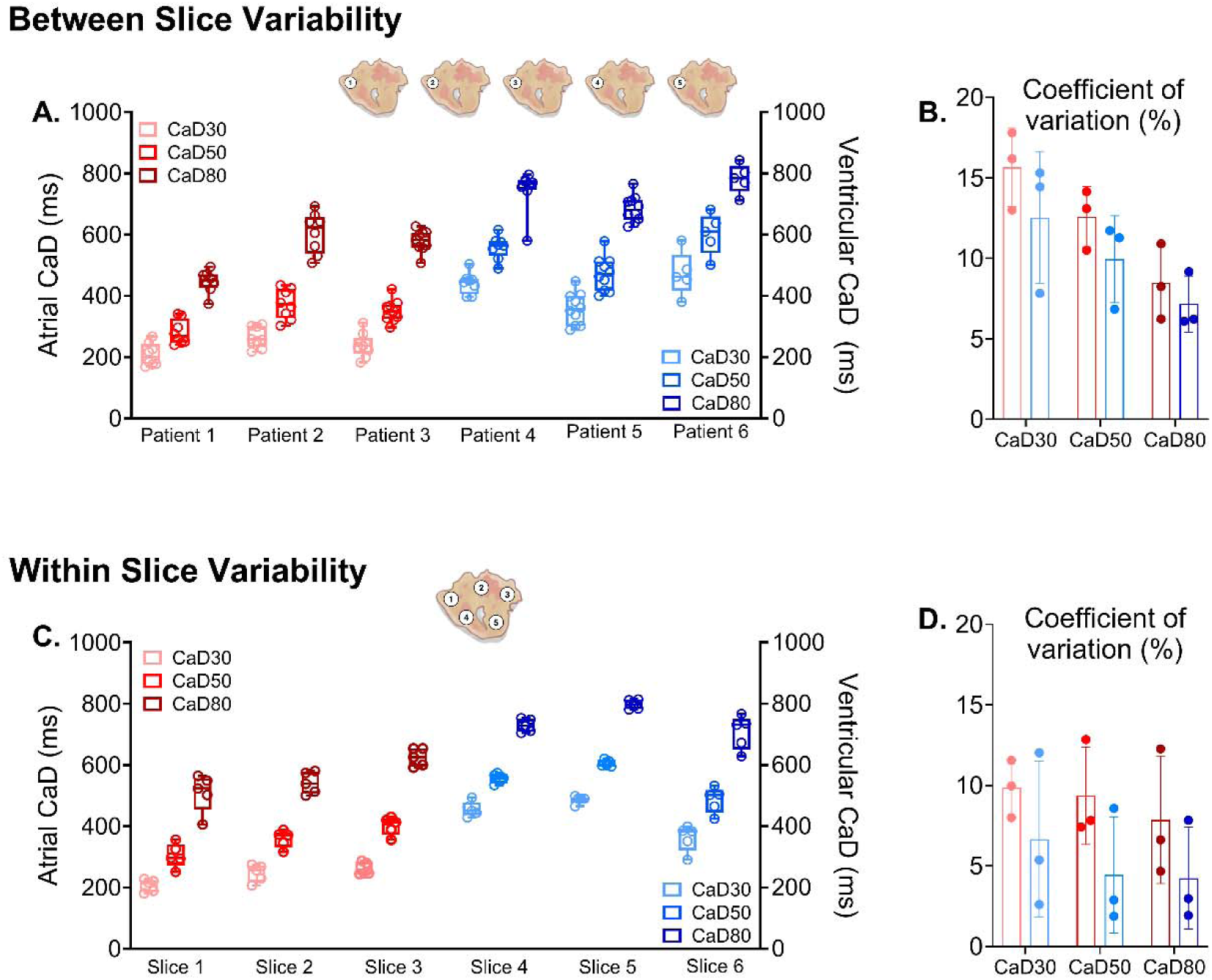
Intra- and interslice variability in calcium transient measurements. **A)** Interslice variability in calcium transient duration was assessed using 5–10 slices per patient. All slices from the same patient were imaged on the same day. **B)** Interslice coefficient of variation. n=6 slices, N=4 patients **C)** Intraslice variability was assessed by measuring calcium transient durations at five distinct regions within each slice. **D)** Intraslice coefficient of variation. n=6 slices, N=6 patients. Calcium transient duration was quantified at 30% (CaD30), 50% (CaD50), and 80% (CaD80) recovery. All recordings were acquired at room temperature using electrical field stimulation at 1 Hz. Atrial preparations are shown in shades of red and ventricular preparations in shades of blue. Calcium transient measurements are shown with minimum– maximum ranges, coefficient of variation data reported as mean ± SD.

### Beta-adrenergic Responsiveness of Human Myocardial Slices

To assess pharmacological responsiveness, atrial and ventricular myocardial slices were treated with isoproterenol or norepinephrine (1 µM; **Figure 4, Supplemental Figure S2**). Previous studies have demonstrated accelerated intracellular calcium-transient kinetics following beta-adrenergic stimulation in human atrial trabeculae and ventricular myocardial slice (Brixius et al., 1997; Kang et al., 2016; Sun et al., 2024). Consistent with these findings, both agents significantly shortened calcium transient durations in atrial and ventricular preparations. Acute isoproterenol exposure significantly shortened CaD30, CaD50, and CaD80 at pacing frequencies of 0.3 and 1 Hz (p < 0.001). At 0.3 Hz, atrial CaD80 decreased from 795.9 ± 289.5 ms at baseline to 538.7 ± 175.9 ms following isoproterenol treatment (−32.3%; p < 0.0001; **Figure 4A**). Similarly, ventricular CaD80 decreased from 1491.4 ± 314.9 ms to 1117.0 ± 357.3 ms (−25.1%; p < 0.0001; **Figure 4C**). Significant shortening was also observed for CaD30 and CaD50 in both chambers, with reductions ranging from 29.8–32.4% in atrial slices and 13.8– 18.8% in ventricular slices.

**Figure 4:**
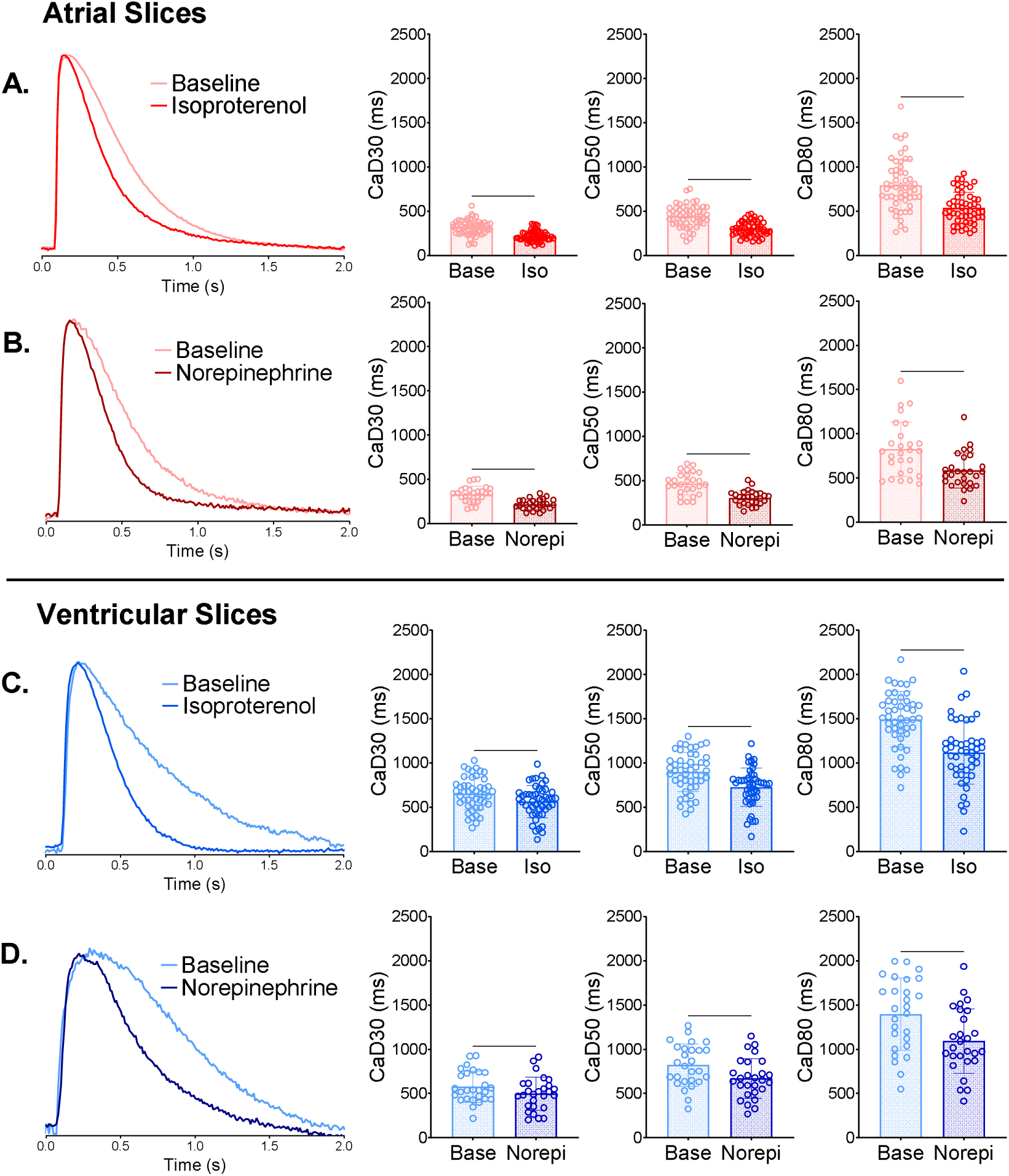
Beta-adrenergic stimulation shortens calcium transient duration in human atrial and ventricular myocardial slices. **A)** Representative intracellular calcium transient recordings and corresponding quantitative analyses from atrial myocardial slices at baseline and following acute exposure to 1 µM isoproterenol (Iso) or 1 µM norepinephrine (Norepi). **(B)** Corresponding calcium transient recordings and analyses from ventricular myocardial slices. Both beta-adrenergic agonists significantly shortened calcium transient duration at 30% (CaD30), 50% (CaD50), and 80% (CaD80) recovery. All recordings were acquired at room temperature during electrical field stimulation at 0.3 Hz. Baseline and post-treatment measurements were obtained from the same myocardial slices. Statistical comparisons were performed using two-tailed, paired Student’s t-tests. Data are presented as mean ± SD. Statistical significance is indicated as **p < 0.01, ***p < 0.001, and ****p < 0.0001. Atrial samples: n=54 slices from N=14 patients; ventricular samples: n=60 slices from N=16 patients. Base = baseline; Iso = isoproterenol; Norepi = norepinephrine.

Norepinephrine exposure produced comparable effects, significantly shortening CaD30, CaD50, and CaD80 in atrial and ventricular preparations paced at 0.3 Hz (p < 0.01). Atrial CaD80 decreased from 824.8 ± 308.0 ms to 584.8 ± 198.3 ms (−29.1%; p < 0.0001; **Figure 4B**), while ventricular CaD80 decreased from 1396.1 ± 409.1 ms to 1091.6 ± 365.9 ms (−21.8%; p < 0.0001; **Figure 4D**). Similar reductions were observed for CaD30 and CaD50, ranging from 34.5–34.6% in atrial slices and 13.7–18.4% in ventricular slices. For both agonists, atrial preparations exhibited greater relative shortening of calcium transient durations than ventricular preparations across the pacing frequencies examined. Collectively, these results demonstrate that human myocardial slices retain functional beta-adrenergic responsiveness, supporting their use for controlled pharmacological assessments.

### Long-Term Preservation of Calcium Cycling and Pharmacological Responsiveness

Given the limited availability of pediatric myocardial specimens, we sought to extend the duration of tissue culture to enable repeated physiological and pharmacological measurements. Calcium transients were recorded during early culture (days 0–5) and subsequently at approximately 10-day intervals using three-day recording windows (days 8–10, 18–20, 28–30, etc.). Longitudinal assessments included 77 atrial slices from seven patients (**Figure 5**) and 43 ventricular slices from seven patients (**Figure 6**). Functional calcium transients were recorded in atrial slices for up to 178 days and ventricular slices for up to 119 days following tissue preparation. These represent the maximum observed culture durations with reliable calcium imaging; culture duration for individual patient specimens is provided in **Table 1**.

**Figure 5:**
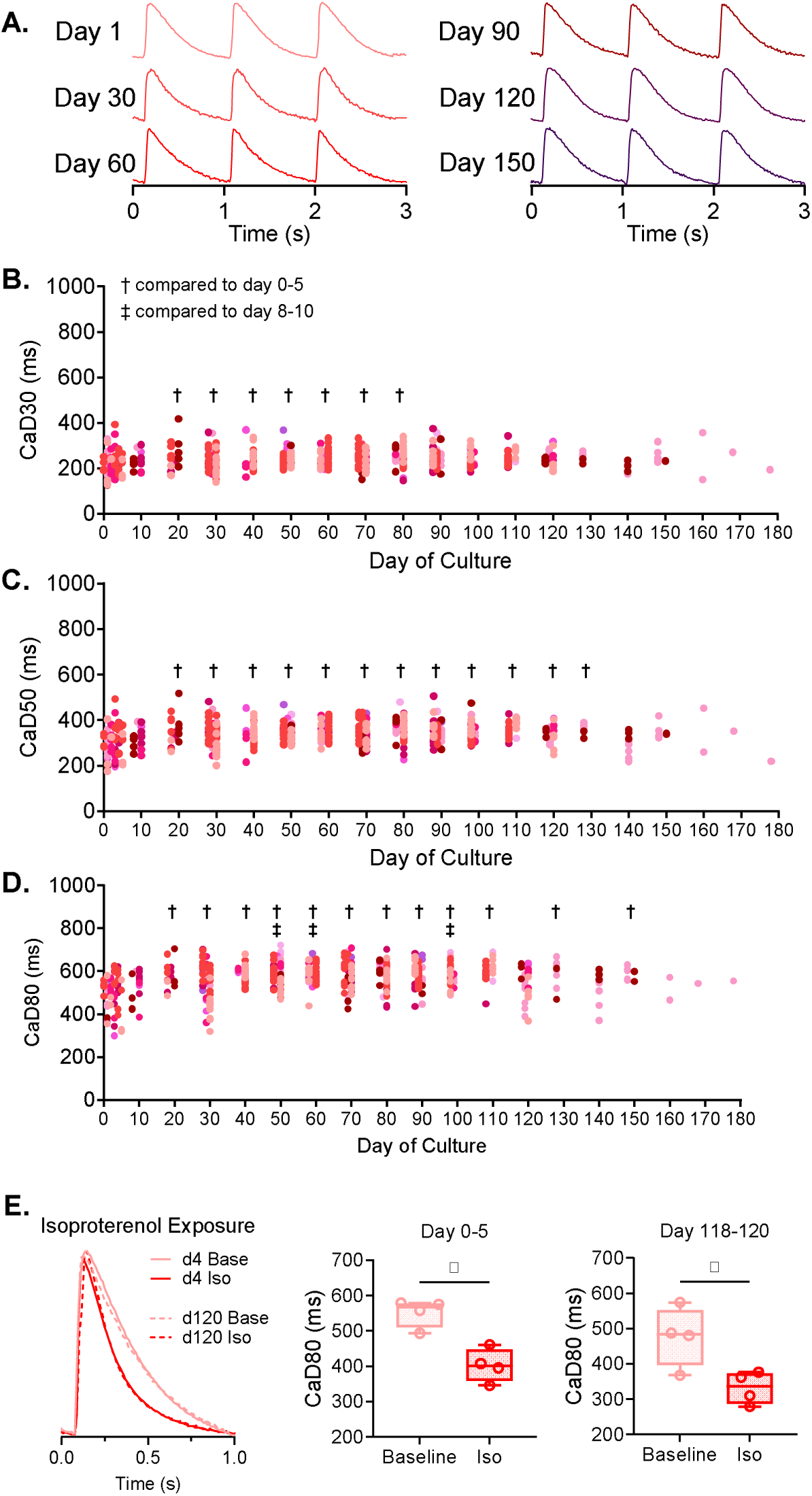
Long-term preservation of calcium cycling and beta-adrenergic responsiveness in human atrial myocardial slices. **A)** Representative intracellular calcium transient recordings obtained at multiple time points during extended culture from right atrial slices collected from a 29-day-old neonate. **B–D)** Longitudinal measurements of calcium transient duration at 30% (CaD30), 50% (CaD50), and 80% (CaD80) recovery over approximately 180 days of passive culture. Data derived from n=77 atrial slices from N=7 individual patients (represented by different shades of red). Statistical comparisons were performed using one-way ANOVA with Holm–Šidák correction for multiple comparisons. †p < 0.05 versus early culture (days 0–5); ‡p < 0.05 versus days 8–10. Only significant comparisons are indicated. **E)** Isoproterenol-induced shortening of CaD80 during early culture (days 0–5; solid line) and extended culture (days 118– 120; dotted line). Paired baseline and post-treatment measurements were obtained from the same slices following acute exposure to 1 µM isoproterenol. Individual slice measurements are shown with minimum–maximum ranges. Statistical comparisons were performed using paired Student’s t-tests; *p < 0.05. Base = baseline; Iso = isoproterenol.

**Figure 6:**
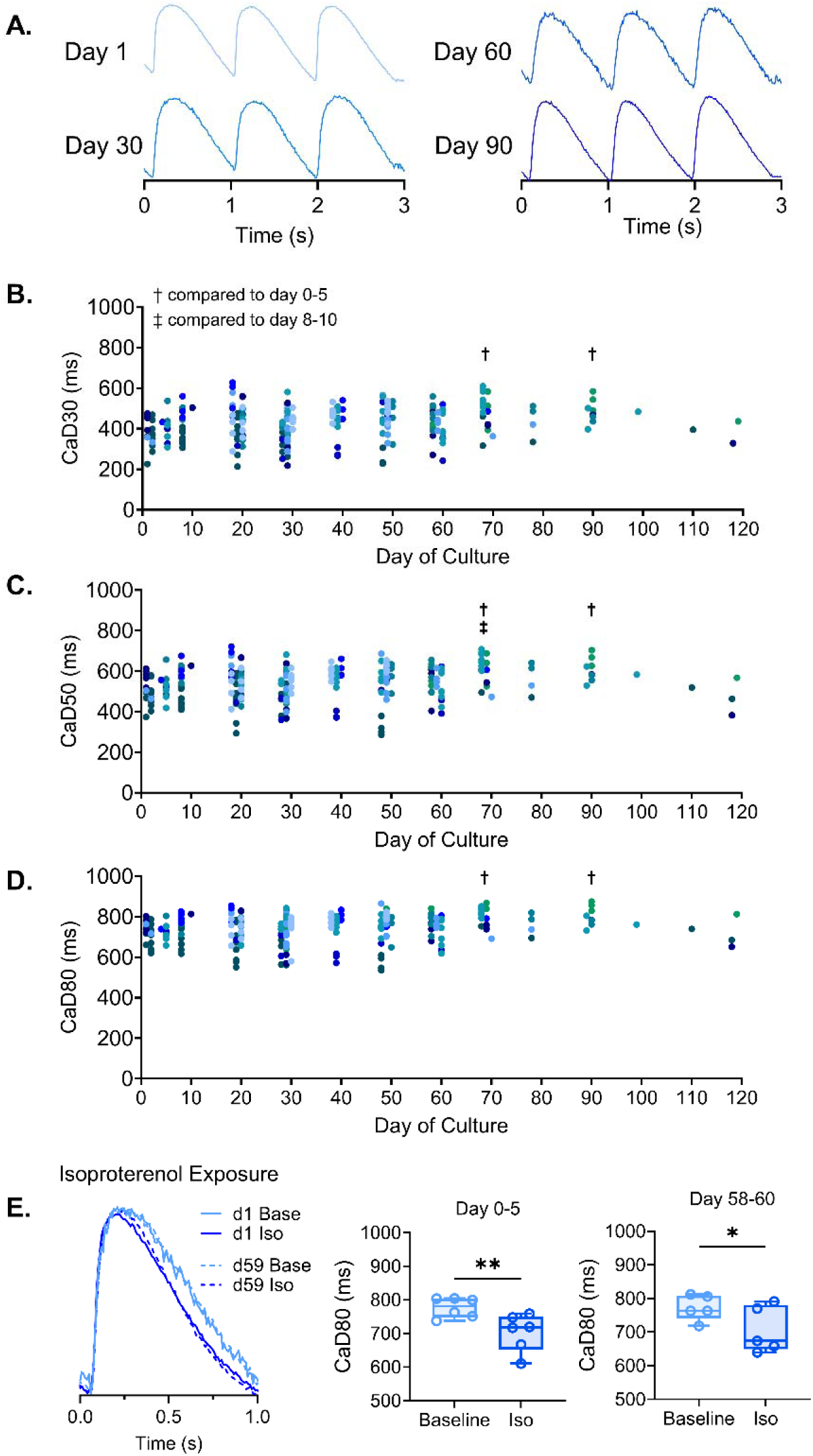
Long-term preservation of calcium cycling and beta-adrenergic responsiveness in human ventricular myocardial slices. **A)** Representative intracellular calcium transient recordings obtained at multiple time points during extended culture from ventricular slices collected from a 2-year-old patient. **B–D)** Longitudinal measurements of calcium transient duration at 30% (CaD30), 50% (CaD50), and 80% (CaD80) recovery over approximately 120 days of passive culture. Data derived from n=43 ventricular slices from N=7 individual patients (represented by different shades of blue). Statistical comparisons were performed using one-way ANOVA with Holm–Šidák correction for multiple comparisons. †p < 0.05 versus early culture (days 0–5); ‡p < 0.05 versus days 8–10. Only significant comparisons are indicated. **E)** Isoproterenol-induced shortening of CaD80 during early culture (days 0–5; solid line) and extended culture (days 58-60; dotted line). Paired baseline and post-treatment measurements were obtained from the same slices following acute exposure to 1 µM isoproterenol. Individual slice measurements are shown with minimum–maximum ranges. Statistical comparisons were performed using paired Student’s t-tests; *p < 0.05. Base = baseline; Iso = isoproterenol.

During the initial weeks of culture, atrial preparations exhibited a modest prolongation of calcium transient duration. Mean CaD80 increased from 495.8 ± 85.8 ms on days 0–5 to 528.2 ± 74.6 ms on days 8–10 and 577.9 ± 56.5 ms on days 18–20 (**Figure 5**). Compared with early culture (days 0–5), CaD80 was significantly prolonged at 12 of 18 subsequent time points (p < 0.01). However, when compared with days 8–10, only three subsequent time points differed significantly (p < 0.05), suggesting an initial period of adaptation(Fischer et al., 2019; Klumm et al., 2022; van der Geest et al., 2025) followed by relatively stable calcium cycling. Ventricular preparations exhibited less variation in calcium transient duration during culture. Mean CaD80 was 722.2 ± 47.5 ms on days 0–5, 742.2 ± 67.1 ms on days 8–10, and 734.7 ± 73.8 ms on days 18–20 (**Figure 6**). No subsequent time points differed significantly from days 8–10, while only two time points differed from the initial days 0–5 measurements (p < 0.05). These findings demonstrate prolonged preservation of functional calcium cycling in both atrial and ventricular myocardial slices, with relatively modest changes following the initial culture period.

We next examined whether pharmacological responsiveness was retained during extended culture by comparing isoproterenol-induced changes in calcium transient duration at early and later time points (**Figure 5,6**: 0.3 Hz data shown). In atrial preparations, 1 µM isoproterenol reduced CaD80 by 26.9% during early culture (days 0–5; 554.6 ± 42.4 to 405.3 ± 47.8 ms; p < 0.05) and by 30.4% on days 118–120 (477.5 ± 84.5 to 332.2 ± 45.6 ms; p < 0.05). Similarly, ventricular preparations exhibited a 9.3% reduction in CaD80 during early culture (772.1 ± 30.6 to 700.3 ± 61.5 ms; p < 0.05) and an 8.5% reduction on days 58–60 (772.6 ± 38.0 to 706.7 ± 68.9 ms; p < 0.05). Although limited to selected preparations, these observations demonstrate that beta-adrenergic responsiveness can be retained over several months of culture, supporting the feasibility of repeated pharmacological assessments using individual human myocardial specimens.

### Extracellular Matrix, Apoptosis-Related Gene Expression, and Tissue Architecture During Culture

To complement our functional assessments, we examined selected genes associated with extracellular matrix remodeling and apoptosis, along with changes in tissue architecture during extended culture (**Figure 7**) (Fischer et al., 2019; Nunez-Toldra et al., 2022). In atrial slices, expression of COL1A1, COL1A2, COL3A1, VIM, and FN1 did not differ significantly across culture compared with days 8–10. Changes in LAMC1 were detected on days 88–90 and 118– 120 (p < 0.05), while POSTN differed only on days 28–30 (p < 0.05). Similarly, most apoptosis-associated transcripts (DFFA, HTRA2, CASP8, CASP9, TP53, BCL2L12, and BOK) showed no significant differences across culture, with a change in CASP10 detected on days 28–30 (p < 0.01). In ventricular slices, COL1A1, COL1A2, and POSTN expression did not differ significantly between days 8–10 and subsequent time points through day 60. Changes in COL3A1 were detected on days 28–30 (p < 0.01), while VIM, LAMC1, and FN1 differed on days 58–60. Among apoptosis-associated transcripts, CASP9, BCL2L12, and BOK showed no significant differences, whereas DFFA and CASP8 differed on days 28–30 and HTRA2 and TP53 on days 58–60 (p < 0.05). CASP10 expression decreased at both later time points (p < 0.001).

**Figure 7:**
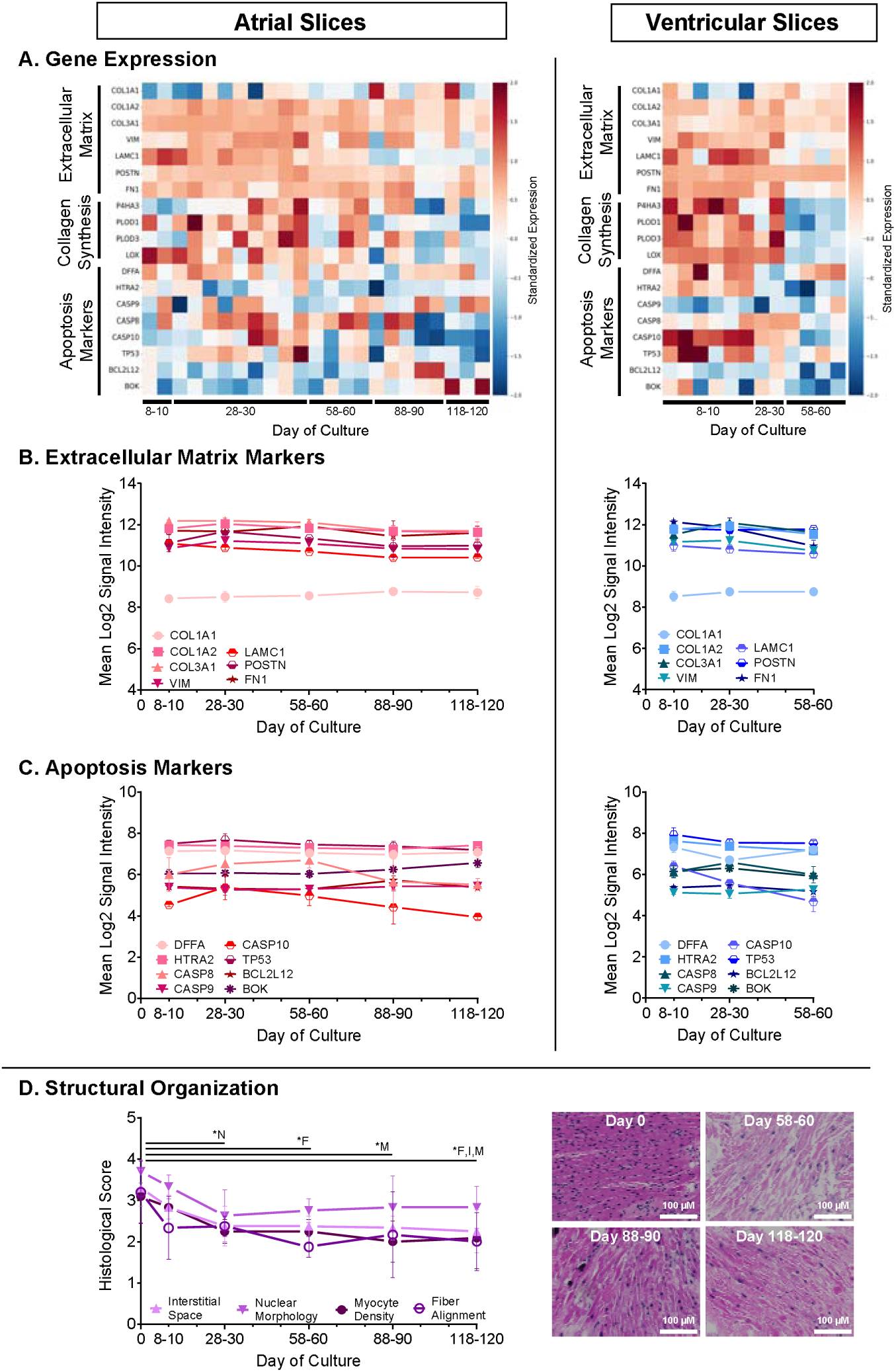
Molecular and histological assessments during extended myocardial slice culture. **A)** Heat maps showing relative expression of selected genes associated with extracellular matrix organization, collagen biosynthesis, and apoptosis-related signaling in atrial and ventricular myocardial slices across culture. Expression values were z-score standardized separately for each gene. **B–C)** Longitudinal measurements of normalized, log₂-transformed microarray signal intensity for selected extracellular matrix-associated and apoptosis-associated transcripts in atrial (red) and ventricular (blue) preparations. **D)** Histological assessment of myocardial tissue architecture during extended culture. Fiber alignment, interstitial space, myocyte density, and nuclear morphology were evaluated using a scoring system (0–4). Horizontal bars indicate significant differences relative to freshly prepared slices (days 0–1; *p < 0.05), with letters identifying the affected parameter: I, interstitial space; M, myocyte density; F, fiber alignment. No significant differences were detected between days 28–30 and subsequent culture time points. Representative hematoxylin and eosin (H&E) stained tissue sections are shown at selected culture time points. Data are presented as mean ± SD. Sample sizes: gene expression, n=48 samples from N=22 patients; histology, n=3 sections from N=15 patients. Scale bars = 100 μm.

We also assessed myocardial architecture using histological scoring of fiber alignment, interstitial space, myocyte density, and nuclear morphology (**Figure 7D**). Compared with freshly prepared slices (days 0–1), significant changes were observed in interstitial space and myocyte density by days 8–10, with differences in myocyte density and fiber alignment persisting at selected later time points (p < 0.05). However, no significant differences were detected in any histological parameter when subsequent time points were compared with days 28–30, suggesting an initial period of structural adaptation followed by relatively stable tissue organization. Together, these exploratory molecular and histological assessments support the preservation of several tissue features during extended culture, although changes in selected extracellular matrix-and apoptosis-associated transcripts were observed.

## DISCUSSION

In this study, we adapted established myocardial slicing methods to develop a practical approach for functional assessment and extended culture of human pediatric cardiac tissue. Using specimens collected during cardiac surgery, we obtained intracellular calcium recordings from preparations representing all four cardiac chambers. Slices exhibited frequency-dependent shortening of calcium transient durations, relatively low intra- and interslice variability, and expected responses to beta-adrenergic stimulation. Importantly, functional calcium transients remained detectable for up to 178 days in atrial slices and 119 days in ventricular slices under passive culture conditions, without continuous electrical stimulation or mechanical loading. These findings demonstrate the feasibility of prolonged physiological assessments using small human myocardial specimens that would otherwise be discarded.

### Functional Validation of Human Myocardial Slices

Previous studies have established that living human myocardial slices retain important aspects of native cardiac physiology, including electrical propagation, calcium cycling, and pharmacological responsiveness(Kang et al., 2016; Qiao et al., 2019). Consistent with these reports, our preparations exhibited the expected frequency-dependent changes in calcium transient duration across all four chambers. Atrial preparations also exhibited shorter calcium transient durations than ventricular preparations, consistent with established differences in human myocardial calcium handling(Kang et al., 2016; Maier et al., 2000). The relatively low intra-and interslice variability further supports the use of this preparation for functional measurements, despite differences in anatomical sampling location and underlying cardiac disease. Although these responses do not establish complete preservation of native cardiac physiology, they demonstrate that small surgical specimens can yield electrically responsive myocardial preparations suitable for quantitative calcium imaging.

### Extended Culture of Human Myocardial Slices

A major advance of this study is the prolonged preservation of functional calcium cycling under relatively simple culture conditions. Previous passive culture protocols have generally supported physiological measurements over several days to weeks (Kang et al., 2016; Ou et al., 2020; Watson et al., 2017). More recently, Zhou et al. demonstrated the feasibility of preparing miniaturized human myocardial slices from small surgical specimens, with reliable contractile function during the first 5–6 days and declining function thereafter(Zhou et al., 2025). Longer culture durations have largely been achieved using specialized biomimetic systems incorporating continuous electrical stimulation and mechanical loading. For example, Fischer et al.(Fischer et al., 2019) demonstrated preservation of contractile function in adult human myocardial slices for up to four months with electromechanical stimulation. In contrast, our preparations retained detectable calcium cycling for approximately six months in atrial tissue and four months in ventricular tissue without continuous electromechanical stimulation. While these studies assess different physiological endpoints and are not directly comparable measures of tissue viability, our findings extend the feasibility of longitudinal calcium imaging using passively cultured human myocardium.

Our culture method incorporated free-floating myocardial slices maintained in continuously agitated medium. Similar approaches using orbital shakers or rocking platforms have been described for human myocardial slices(George et al., 2020; Qiao et al., 2019), while other studies have demonstrated that medium agitation can influence the preservation of myocardial function(Cao-Ehlker et al., 2023; Fischer et al., 2019). In our preparations, continuous rocking maintained slices submerged in culture medium without requiring specialized mechanical loading devices. Although we did not systematically compare culture conditions, this approach provides an accessible method for maintaining functional myocardial preparations over extended periods.

### Functional Adaptation during Extended Culture

We observed modest changes in calcium transient duration during the initial weeks of culture, followed by relatively stable measurements at later time points. Similar periods of functional adaptation have been reported in previous myocardial tissue culture studies. Fischer et al. (Fischer et al., 2019) observed an initial decline in contractile force followed by gradual recovery during prolonged culture, while Watson et al.(Watson et al., 2017) and Klumm et al.(Klumm et al., 2022) documented early changes in contractile performance in animal myocardial slices and human atrial trabeculae, respectively. More recently, van der Geest et al. (van der Geest et al., 2025) reported improved calcium transient amplitude, pacing responsiveness, and calcium reuptake kinetics following approximately 10 days of human myocardial slice culture, alongside other functional and molecular changes. These findings suggest that myocardial preparations can undergo substantial physiological adjustment during early culture, although the direction and magnitude of these changes may depend on experimental conditions. Interestingly, we occasionally observed myocardial preparations that initially lacked detectable calcium transients or contractile activity but subsequently became responsive during the first several days of culture. This qualitative observation may reflect recovery from the tissue slicing procedure and/or adaptation to culture conditions, although the underlying mechanisms remain uncertain. Collectively, our findings and previous reports suggest that an initial equilibration period may be beneficial before conducting longitudinal or repeated pharmacological assessments.

### Preservation of Pharmacological Responsiveness

Our results also demonstrate that beta-adrenergic responsiveness is retained in human myocardial slices. Both isoproterenol and norepinephrine shortened calcium transient durations in atrial and ventricular preparations, consistent with established effects of beta-adrenergic stimulation on intracellular calcium cycling. Similar responses have been demonstrated in isolated human atrial trabeculae(Brixius et al., 1997) and adult human myocardial slices(Kang et al., 2016; Sun et al., 2024). Importantly, we observed comparable directional responses to isoproterenol during early and extended culture, with significant shortening of CaD80 at approximately four months in atrial preparations and two months in ventricular preparations. Although these observations were limited to selected specimens and do not establish equivalent pharmacological sensitivity across time, they demonstrate that measurable drug responsiveness can persist throughout extended culture. This provides an opportunity to obtain repeated measurements from individual surgical specimens and potentially investigate the consequences of longer-term exposures.

### Molecular and Structural Assessments during Extended Culture

Functional preservation does not necessarily indicate that myocardial structure or gene expression remains unchanged. Previous studies have demonstrated culture-associated remodeling, including changes in extracellular matrix organization, cytoskeletal pathways, and cellular metabolism(Fischer et al., 2019; Nunez-Toldra et al., 2022; van der Geest et al., 2025). Notably, van der Geest et al.(van der Geest et al., 2025) observed preservation of sarcomeric and mitochondrial organization despite substantial transcriptional changes following 10 days of culture. In our study, exploratory assessments of selected extracellular matrix-and apoptosis-associated transcripts revealed relatively limited changes in several markers, although individual genes differed at later time points. Histological assessments similarly demonstrated preservation of several architectural features following the initial culture period. These findings provide complementary evidence of tissue preservation but do not exclude broader molecular remodeling during extended culture.

### Potential Applications

Human pediatric myocardial specimens are an underutilized resource for investigating developmental cardiac physiology and pharmacology. This is particularly relevant to children with CHD, who undergo substantial postnatal myocardial maturation while frequently receiving cardiovascular medications with limited pediatric-specific safety and efficacy data(Pasquali et al., 2008). Our approach permits direct physiological measurements using myocardium from patients across a broad developmental age range and with diverse cardiac diagnoses. Extending the functional lifespan of these preparations may also increase the number of experiments performed using individual surgical specimens. Future applications could incorporate optical action potential measurements, microelectrode arrays, and direct assessments of contractility to expand the range of physiological and pharmacological endpoints. Overall, this study establishes a practical approach for repeated functional assessment of human myocardial tissue, supporting future investigations of developmental and disease-associated cardiac physiology.

### Study Limitations

Our study has several limitations that should be considered. First, specimens were primarily obtained from patients with CHD or other cardiac conditions, and their physiological properties may not represent healthy pediatric myocardium. Patient ages, diagnoses, and anatomical sampling locations were heterogeneous, and some chamber-specific comparisons were limited by specimen availability. In particular, left atrial recordings originated from a single adult patient. Second, calcium imaging was performed at room temperature, which can alter calcium transient kinetics(Chung & Campbell, 2013). Third, extended culture was evaluated in a limited number of donors, with multiple slices derived from individual specimens. Maximum recording durations therefore do not represent population-level survival or functional preservation rates. Viability measurements were also limited to a subset of acute preparations. Fourth, our physiological assessments focused on calcium cycling and selected beta-adrenergic responses; direct measurements of membrane potential and contractile force were not performed. Finally, molecular and histological assessments were exploratory, and the contributions of individual culture conditions were not independently evaluated. Additional studies with larger donor cohorts and complementary physiological endpoints will be needed to establish the broader applicability and reproducibility of this approach.

## Acknowledgements

We acknowledge members of the cardiac surgery operating room team, including Aybala Tongut, Blezzy Bote, Alecia Byrd, Jannette Callos, Kenisha Cyrus, Moozhda Hanif, Evelyn Ravizee, Sandra Sunderland, and Hyung Mi (Grace) Yang for assistance with biospecimen collection. We gratefully acknowledge the Children’s National Research Institute Genomics & Bioinformatics Core, including Susan Knoblach and Dan Tran, for assistance with gene expression experiments. We thank the Pathology & Laboratory Medicine Research Core at Children’s National Hospital for help with paraffin embedding, slicing of formalin fixed samples, and H&E staining. Microscope analysis was carried out at the Children’s National Research Institute (CNRI) Light Microscopy and Image Analysis Core supported by CNRI and NIH grant P30HD040677.

## Author Contributions

DG and NGP designed and conceived experiments; DG, AB, MD, and YD procured biospecimens; DG, KM, AF, JP, and NGP performed experiments; DG, KM, AF, JP, and NGP analyzed data; DG, KM, AF, JP, and NGP interpreted results of experiments; DG, KM, AF, and NGP prepared figures; DG and NGP drafted manuscript; all authors edited and approved the final manuscript.

## Supplemental File

**Supplemental Figure S1.**
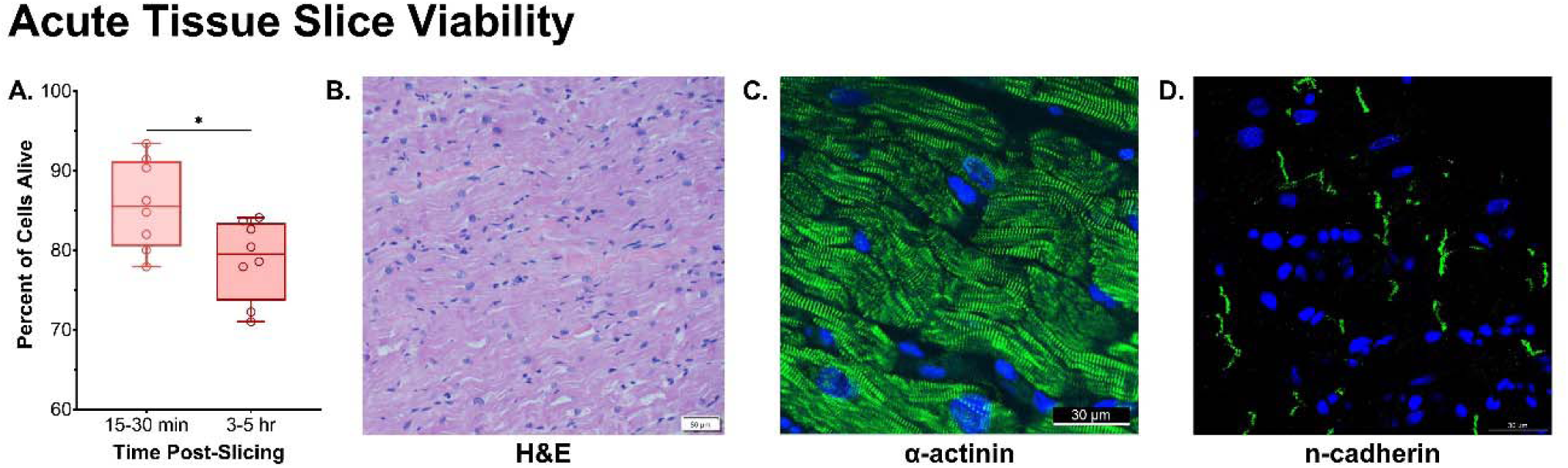
Acute cell viability and representative histological features of human myocardial slices. **A)** Percentage of viable cells at 15–30 minutes and 3–5 hours following tissue preparation. Individual measurements are shown with minimum–maximum ranges. Student’s t-test, *p < 0.05. **B–D)** Representative images of freshly prepared myocardial slices (day 0), showing **B)** hematoxylin and eosin (H&E) staining, **C)** sarcomeric α-actinin (green) and nuclei (DAPI, blue), and **D)** N-cadherin (green) and nuclei (DAPI, blue). Images illustrate preservation of recognizable myocardial architecture, including sarcomeric organization and intercellular junction-associated protein localization. Scale bars are indicated in each panel.

**Supplemental Figure 2:**
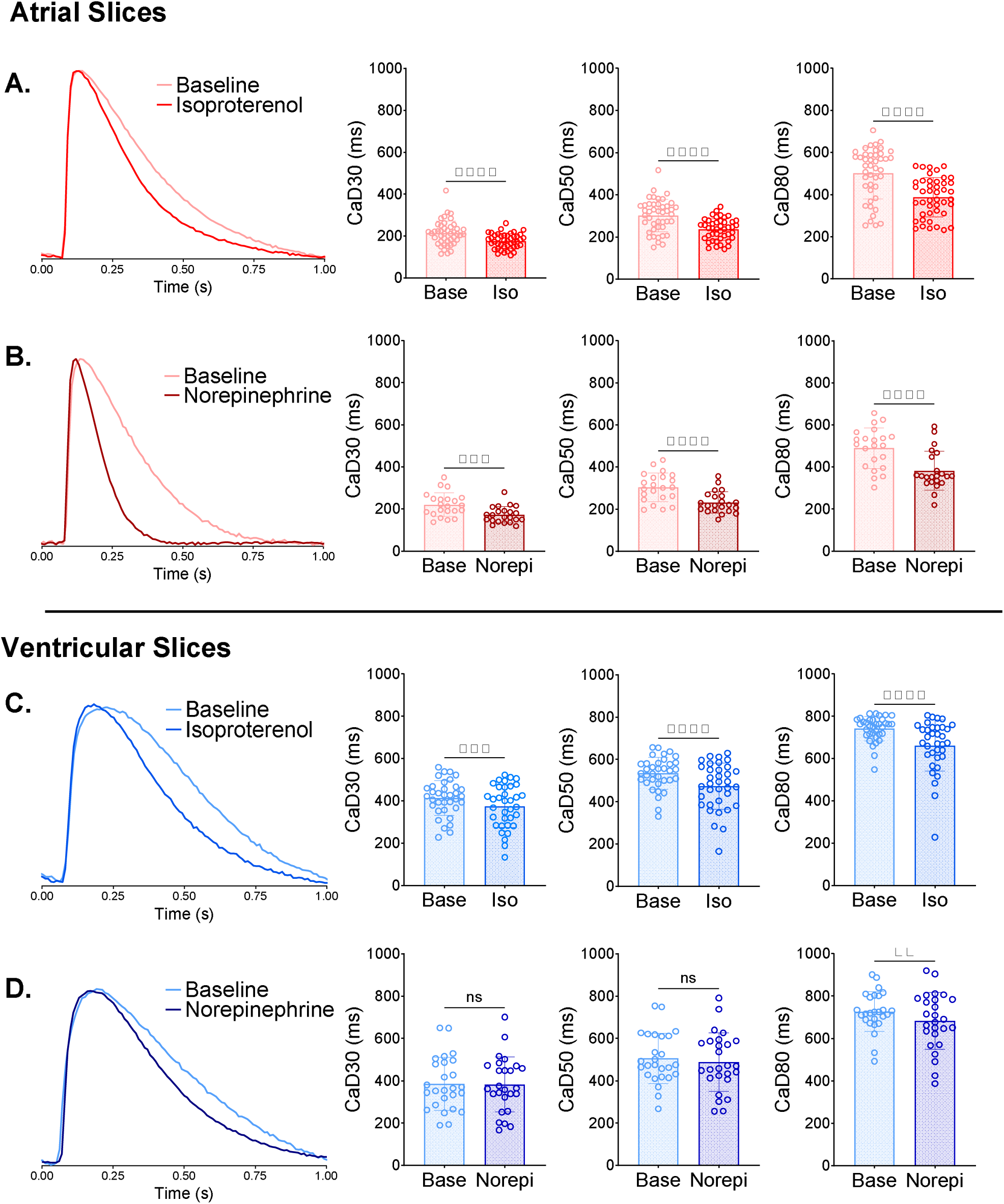
Beta-adrenergic stimulation shortens calcium transient duration in human atrial and ventricular myocardial slices (1 Hz) **A)** Representative intracellular calcium transient recordings and corresponding quantitative analyses from atrial myocardial slices at baseline and following acute exposure to 1 µM isoproterenol (Iso) or **B)** 1 µM norepinephrine (Norepi). **C-D)** Corresponding calcium transient recordings and analyses from ventricular myocardial slices. All recordings were acquired at room temperature during electrical field stimulation at 1 Hz. Baseline and post-treatment measurements were obtained from the same myocardial slices. Statistical comparisons were performed using two-tailed, paired Student’s t-tests. Data are presented as mean ± SD. Statistical significance is indicated as **p < 0.01, ***p < 0.001, and ****p < 0.0001. Atrial samples: n=54 slices from N=14 patients; ventricular samples: n=60 slices from N=16 patients. Base = baseline; Iso = isoproterenol; Norepi = norepinephrine.

## Supplemental Figures

**Supplemental Table S1:** Tyrode’s Slicing Solution.

| Chemical | Concentration | Volume | Catalog # | Supplier |
|---|---|---|---|---|
| Deionized Water | - | 1 L | - | - |
| D-Glucose Anhydrous | 1.0 mM | 1.86 g | 0188-500G | VWR Life Science |
| HEPES | 10.0 mM | 2.38 g | H3375-500G | Sigma-Aldrich |
| Potassium Chloride | 4.5 mM | 0.335 g | P3911-1KG | Sigma-Aldrich |
| Sodium Chloride | 140.0 mM | 8.18 g | S9888-5KG | Sigma-Aldrich |
| Magnesium Chloride, 1M | 1.0 mM | 1 mL | E525-500ML | VWR Life Science |
| Calcium Chloride, 1M | 1.8 mM | 1.8 mL | E506-500ML | VWR Life Science |
| 2,3-Butanedione Monoxime (BDM) | 30 mM | 3.0 g | B0753-100G | Sigma-Aldrich |
| * pH to 7.3 |  |  |  |  |

**Supplemental Table S2:** Tyrode’s Recovery Solution.

| Chemical | Concentration | Volume | Catalog # | Supplier |
|---|---|---|---|---|
| Deionized Water | - | 1 L | - | - |
| D-Glucose Anhydrous | 1.0 mM | 1.86 g | 0188-500G | VWR Life Science |
| HEPES | 10.0 mM | 2.38 g | H3375-500G | Sigma-Aldrich |
| Potassium Chloride | 4.5 mM | 0.335 g | P3911-1KG | Sigma-Aldrich |
| Sodium Chloride | 140.0 mM | 8.18 g | S9888-5KG | Sigma-Aldrich |
| Magnesium Chloride, 1M | 1.0 mM | 1 mL | E525-500ML | VWR Life Science |
| Calcium Chloride, 1M | 1.8 mM | 1.8 mL | E506-500ML | VWR Life Science |
| * pH to 7.3 |  |  |  |  |

**Supplemental Table S3:**
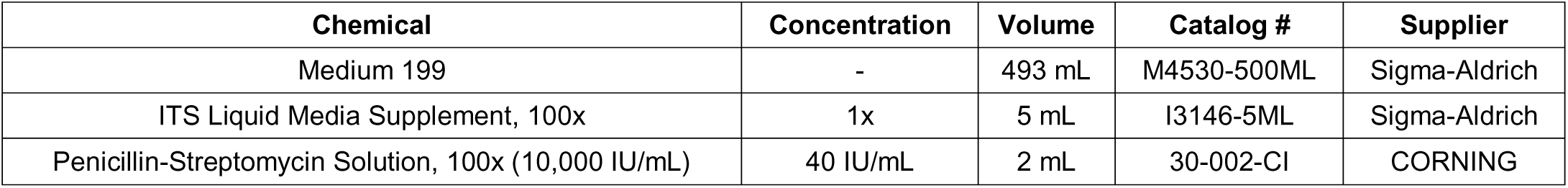
M199 Culture Solution.

## References

Blackwell, D. J., Schmeckpeper, J., Blackwell, D. J., Schmeckpeper, J., & Knollmann, B. C. (2022). Animal Models to Study Cardiac Arrhythmias. Circulation Research, 130(12), 1926–1964. 10.1161/CIRCRESAHA.122.320258

Brandenburger, M., Wenzel, J., Bogdan, R., Richardt, D., Nguemo, F., Reppel, M., Hescheler, J., Terlau, H., & Dendorfer, A. (2012). Organotypic slice culture from human adult ventricular myocardium. Cardiovascular Research, 93(1), 50–59. 10.1093/CVR/CVR259

Brixius, K., Pietsch, M., Hoischen, S., Müller-Ehmsen, J., & Schwinger, R. H. G. (1997). Effect of inotropic interventions on contraction and Ca2+ transients in the human heart. Https://Doi.Org/10.1152/Jappl.1997.83.2.652, 83(2), 652–660. 10.1152/JAPPL.1997.83.2.652

Cao-Ehlker, X., Fischer, C., Lu, K., Bruegmann, T., Sasse, P., Dendorfer, A., & Tomasi, R. (2023). Optimized Conditions for the Long-Term Maintenance of Precision-Cut Murine Myocardium in Biomimetic Tissue Culture. Bioengineering, 10(2), 171. 10.3390/BIOENGINEERING10020171

Chen, E. Y., Tan, C. M., Kou, Y., Duan, Q., Wang, Z., Meirelles, G. V., Clark, N. R., & Ma’ayan, A. (2013). Enrichr: interactive and collaborative HTML5 gene list enrichment analysis tool. BMC Bioinformatics, 14. 10.1186/1471-2105-14-128

Chung, C. S., & Campbell, K. S. (2013). Temperature and transmural region influence functional measurements in unloaded left ventricular cardiomyocytes. Physiological Reports, 1(6), e00158–e00158. 10.1002/phy2.158

Elhamine, F., Iorga, B., Krüger, M., Hunger, M., Eckhardt, J., Sreeram, N., Bennink, G., Brockmeier, K., Pfitzer, G., & Stehle, R. (2016). Postnatal development of right ventricular myofibrillar biomechanics in relation to the sarcomeric protein phenotype in pediatric patients with conotruncal heart defects. Journal of the American Heart Association, 5(6). 10.1161/JAHA.116.003699

Fischer, C., Milting, H., Fein, E., Reiser, E., Lu, K., Seidel, T., Schinner, C., Schwarzmayr, T., Schramm, R., Tomasi, R., Husse, B., Cao-Ehlker, X., Pohl, U., & Dendorfer, A. (2019). Long-term functional and structural preservation of precision-cut human myocardium under continuous electromechanical stimulation in vitro. Nature Communications, 10(1). 10.1038/S41467-018-08003-1

George, S. A., Brennan, J. A., & Efimov, I. R. (2020). Preclinical Cardiac Electrophysiology Assessment by Dual Voltage and Calcium Optical Mapping of Human Organotypic Cardiac Slices. Journal of Visualized Experiments : JoVE, 2020(160), 1–17. 10.3791/60781

Guo, Y., & Pu, W. T. (2020). Cardiomyocyte Maturation: New Phase in Development. Circulation Research, 126(8), 1086–1106. 10.1161/CIRCRESAHA.119.315862/FORMAT/EPUB

Janssen, P. M. L., Biesiadecki, B. J., Ziolo, M. T., & Davis, J. P. (2016). The Need for Speed; Mice, Men, and Myocardial Kinetic Reserve. Circulation Research, 119(3), 418. 10.1161/CIRCRESAHA.116.309126

Kang, C., Qiao, Y., Li, G., Baechle, K., Camelliti, P., Rentschler, S., & Efimov, I. R. (2016). Human Organotypic Cultured Cardiac Slices: New Platform For High Throughput Preclinical Human Trials. Scientific Reports, 6(1), 28798. 10.1038/srep28798

Karbassi, E., Fenix, A., Marchiano, S., Muraoka, N., Nakamura, K., Yang, X., & Murry, C. E. (2020). Cardiomyocyte maturation: advances in knowledge and implications for regenerative medicine. Nature Reviews. Cardiology, 17(6), 341–359. 10.1038/S41569-019-0331-X

Klumm, M. J., Heim, C., Fiegle, D. J., Weyand, M., Volk, T., & Seidel, T. (2022). Long-Term Cultivation of Human Atrial Myocardium. Frontiers in Physiology, 13, 839139. 10.3389/FPHYS.2022.839139/TEXT

Kuleshov, M. V., Jones, M. R., Rouillard, A. D., Fernandez, N. F., Duan, Q., Wang, Z., Koplev, S., Jenkins, S. L., Jagodnik, K. M., Lachmann, A., McDermott, M. G., Monteiro, C. D., Gundersen, G. W., & Maayan, A. (2016). Enrichr: a comprehensive gene set enrichment analysis web server 2016 update. Nucleic Acids Research, 44(W1), W90–W97. 10.1093/NAR/GKW377

Linscheid, N., Santos, A., Poulsen, P. C., Mills, R. W., Calloe, K., Leurs, U., Ye, J. Z., Stolte, C., Thomsen, M. B., Bentzen, B. H., Lundegaard, P. R., Olesen, M. S., Jensen, L. J., Olsen, J. V., & Lundby, A. (2021). Quantitative proteome comparison of human hearts with those of model organisms. PLoS Biology, 19(4), e3001144. 10.1371/JOURNAL.PBIO.3001144

Lou, Q., Fedorov, V. V., Glukhov, A. V., Moazami, N., Fast, V. G., & Efimov, I. R. (2011). Transmural Heterogeneity and Remodeling of Ventricular Excitation-Contraction Coupling in Human Heart Failure. Circulation, 123(17), 1881. 10.1161/CIRCULATIONAHA.110.989707

Maier, L. S., Barckhausen, P., Weisser, J., Aleksic, I., Baryalei, M., & Pieske, B. (2000). Ca2+ handling in isolated human atrial myocardium. Https://Doi.Org/10.1152/Ajpheart.2000.279.3.H952, 279(3 48-3). 10.1152/AJPHEART.2000.279.3.H952

Mandalenakis, Z., Giang, K. W., Eriksson, P., Liden, H., Synnergren, M., Wåhlander, H., Fedchenko, M., Rosengren, A., & Dellborg, M. (2020). Survival in children with congenital heart disease: Have we reached a peak at 97%? Journal of the American Heart Association, 9(22), 17704. 10.1161/JAHA.120.017704/SUPPL_FILE/JAH35659-SUP-0001-SUPINFO.PDF

Milani-Nejad, N., & Janssen, P. M. L. L. M. L. (2014). Small and large animal models in cardiac contraction research: advantages and disadvantages. Pharmacology & Therapeutics, 141(3), 235–249. 10.1016/j.pharmthera.2013.10.007

Nunez-Toldra, R., Kirwin, T., Ferraro, E., Pitoulis, F. G., Nicastro, L., Bardi, I., Kit-Anan, W., Gorelik, J., Simon, A. R., & Terracciano, C. M. (2022). Mechanosensitive molecular mechanisms of myocardial fibrosis in living myocardial slices. ESC Heart Failure, 9(2), 1400. 10.1002/EHF2.13832

Odening, K. E., Gomez, A. M., Dobrev, D., Fabritz, L., Heinzel, F. R., Mangoni, M. E., Molina, C. E., Sacconi, L., Smith, G., Stengl, M., Thomas, D., Zaza, A., Remme, C. A., & Heijman, J. (2021). ESC working group on cardiac cellular electrophysiology position paper: relevance, opportunities, and limitations of experimental models for cardiac electrophysiology research. *Europace : European Pacing, Arrhythmias, and Cardiac Electrophysiology : Journal of the Working Groups on Cardiac Pacing*, Arrhythmias, and Cardiac Cellular Electrophysiology of the European Society of Cardiology, 23(11), 1795–1814. 10.1093/EUROPACE/EUAB142

Ou, Q., Abouleisa, R. R. E., Tang, X. L., Juhardeen, H. R., Meki, M. H., Miller, J. M., Giridharan, G., El-Baz, A., Bolli, R., & Mohamed, T. M. A. (2020). Slicing and Culturing Pig Hearts under Physiological Conditions. Journal of Visualized Experiments : JoVE, 2020(157). 10.3791/60913

Ou, Q., Jacobson, Z., Abouleisa, R. R. E., Tang, X. L., Hindi, S. M., Kumar, A., Ivey, K. N., Giridharan, G., El-Baz, A., Brittian, K., Rood, B., Lin, Y. H., Watson, S. A., Perbellini, F., McKinsey, T. A., Hill, B. G., Jones, S. P., Terracciano, C. M., Bolli, R., & Mohamed, T. M. A. (2019). Physiological Biomimetic Culture System for Pig and Human Heart Slices. Circulation Research, 125(6), 628–642. 10.1161/CIRCRESAHA.119.314996

Pasquali, S. K., Hall, M., Slonim, A. D., Jenkins, K. J., Marino, B. S., Cohen, M. S., & Shah, S. S. (2008). Off-Label Use of Cardiovascular Medications in Children Hospitalized With Congenital and Acquired Heart Disease. 1(2), 74–83. 10.1161/CIRCOUTCOMES.108.787176

Qiao, Y., Dong, Q., Li, B., Obaid, S., Miccile, C., Yin, R. T., Talapatra, T., Lin, Z., Li, S., Li, Z., & Efimov, I. R. (2019). Multiparametric slice culture platform for the investigation of human cardiac tissue physiology. Progress in Biophysics and Molecular Biology, 144, 139–150. 10.1016/j.pbiomolbio.2018.06.001

Ripplinger, C. M., Glukhov, A. V., Kay, M. W., Boukens, B. J., Chiamvimonvat, N., Delisle, B. P., Fabritz, L., Hund, T. J., Knollmann, B. C. B. C., Li, N., Murray, K. T., Poelzing, S., Quinn, T. A., Remme, C. A., Rentschler, S. L., Rose, R. A., & Posnack, N. G. (2022). Guidelines for assessment of cardiac electrophysiology and arrhythmias in small animals. American Journal of Physiology. Heart and Circulatory Physiology, 323(6), H1137–H1166. https://pubmed.ncbi.nlm.nih.gov/36269644/

Salameh, S., Guerrelli, D., Miller, J. A., Desai, M., Moise, N., Yerebakan, C., Bruce, A., Sinha, P., D’Udekem, Y., Weinberg, S. H., & Posnack, N. G. (2024). Connecting transcriptomics with computational modeling to reveal developmental adaptations in pediatric human atrial tissue. American Journal of Physiology-Heart and Circulatory Physiology, 327(6), H1413– H1430. 10.1152/AJPHEART.00474.2024

Salameh, S., Guerrelli, D., Swift, L. M., Haski, A., Bruce, A., Desai, M., d’Udekem, Y., & Posnack, N. G. (2026). Chamber-specific transcriptomic insight into cardiac development using guinea pig and human heart tissue. Physiological Genomics, 58(1), 1–11. 10.1152/physiolgenomics.00212.2025

Salameh, S., Ogueri, V., & Posnack, N. G. (2023). Adapting to a new environment: postnatal maturation of the human cardiomyocyte. The Journal of Physiology, 601(13), 2593–2619. 10.1113/JP283792

Sun, Z., Lu, K., Kamla, C., Kameritsch, P., Seidel, T., & Dendorfer, A. (2024). Synchronous force and Ca2+ measurements for repeated characterization of excitation-contraction coupling in human myocardium. Communications Biology *2024* 7:*1*, *7*(1), 220-. 10.1038/s42003-024-05886-3

Usta, E., Renovanz, M., Mustafi, M., Ziemer, G., & Aebert, H. (2010). Human cardiac tissue in a microperfusion chamber simulating extracorporeal circulation-ischemia and apoptosis studies. Journal of Cardiothoracic Surgery 2010 5:1, *5*(1), 3-. 10.1186/1749-8090-5-3

Van Den Berg, C. W., Okawa, S., Chuva De Sousa Lopes, S. M., Van Iperen, L., Passier, R., Braam, S. R., Tertoolen, L. G., Del Sol, A., Davis, R. P., & Mummery, C. L. (2015). Transcriptome of human foetal heart compared with cardiomyocytes from pluripotent stem cells. Development (Cambridge, England), 142(18), 3231–3238. 10.1242/DEV.123810

van der Geest, J. S. A., Benavente, E. D., van Ham, W. B., Doevendans, P. A., van Laake, L. W., de Boer, T. P., Sampaio-Pinto, V., & Sluijter, J. P. G. (2025). Characterization of human living myocardial slices culture-induced adaptations: a translational perspective. Journal of Molecular and Cellular Cardiology Plus, 13. 10.1016/j.jmccpl.2025.100465

Van Der Geest, J. S. A., De Boer, T. P., Terracciano, C. M., Thum, T., Dendorfer, A., Doevendans, P. A., Van Laake, L. W., Sluijter, J. P. G., & Sampaio-Pinto, V. (2025). Living myocardial slices: walking the path towards standardization. Cardiovascular Research, 121(7), 1011–1023. 10.1093/CVR/CVAF079

Van Der Linde, D., Konings, E. E. M., Slager, M. A., Witsenburg, M., Helbing, W. A., Takkenberg, J. J. M., & Roos-Hesselink, J. W. (2011). Birth prevalence of congenital heart disease worldwide: A systematic review and meta-analysis. Journal of the American College of Cardiology, 58(21), 2241–2247. 10.1016/j.jacc.2011.08.025

Waleczek, F. J. G., Cipriano, G., Haas, J. A., Garg, A., Pfanne, A., Just, A., Neumüller, S., Hegermann, J., Pich, A., Radocaj, A., Xiao, K., Weber, N., & Thum, T. (2025). Prolonged Hypoxia in Rat Living Myocardial Slices Affects Function, Expression, and Structure. International Journal of Molecular Sciences, 26(1), 218. 10.3390/IJMS26010218/S1

Watson, S. A., Dendorfer, A., Thum, T., & Perbellini, F. (2020). A practical guide for investigating cardiac physiology using living myocardial slices. Basic Research in Cardiology 2020 115:*6*, *115*(6), 61-. 10.1007/S00395-020-00822-Y

Watson, S. A., Scigliano, M., Bardi, I., Ascione, R., Terracciano, C. M., & Perbellini, F. (2017). Preparation of viable adult ventricular myocardial slices from large and small mammals. Nature Protocols, 12(12), 2623–2639. 10.1038/NPROT.2017.139

Wen, Q., Gandhi, K., Capel, R. A., Hao, G., O’Shea, C., Neagu, G., Pearcey, S., Pavlovic, D., Terrar, D. A., Wu, J., Faggian, G., Camelliti, P., & Lei, M. (2018). Transverse cardiac slicing and optical imaging for analysis of transmural gradients in membrane potential and Ca2+ transients in murine heart. The Journal of Physiology, 596(17), 3951–3965. 10.1113/JP276239

Wiegerinck, R. F., Cojoc, A., Zeidenweber, C. M., Ding, G., Shen, M., Joyner, R. W., Fernandez, J. D., Kanter, K. R., Kirshbom, P. M., Kogon, B. E., Wagner, M. B., Rob F. Wiegerinck, Anca Cojoc, Carlo M. Zeidenweber, Guoliang Ding, Ming Shen, Ronald W. Joyner, Janet D. Fernandez, Kirk R. Kanter, … Mary B. Wagner. (2009). Force Frequency Relationship of the Human Ventricle Increases During Early Postnatal Development. Pediatric Re, 65(4), 414–419. 10.1203/PDR.0b013e318199093c

Zhou, Z., van Steenis, Y. P., Henry, S., van Doorn, E. C. H., Amesz, J. H., van de Woestijne, P. C., de Groot, N. M. S., Manintveld, O. C., Bartelds, B., & Taverne, Y. J. H. J. (2025). Viability and Longevity of Human Miniaturized Living Myocardial Slices. Journal of Cardiovascular Development and Disease, 12(7). 10.3390/JCDD12070269

